# Reef fish escape responses selectively match predator attack speeds

**DOI:** 10.64898/2026.03.21.713327

**Authors:** Sara Linde Neven, Lena Faber, Benjamin T. Martin

## Abstract

Animals must continually balance foraging with the risk of predation. In complex natural environments, this means quickly distinguishing between threats and harmless situations. We investigated how site-associated coral reef fishes decide to escape in response to visual cues mimicking predator attacks, using controlled underwater presentations of looming stimuli at varying speeds. We measured escape responses across species and social contexts, comparing them to predator attack speeds observed in the same habitat. Escape responses were highly sensitive to the speed of the looming stimulus, with no responses occurring at low speeds. The speeds triggering escape matched those of predator attacks, whereas cruising swim speeds never triggered a response. Species employed distinct antipredator strategies: Brown Chromis foraged away from shelter with high responsiveness, whereas Bicolor Damselfish remained shelter-dependent with lower escape propensities. Contrary to expectations, the social factors did not affect responses in this study. These findings demonstrate that reef fish are highly sensitive to the approach speed of objects, with species-specific strategies further shaping behaviors. By combining realistic visual threats with natural predator attack data, this study offers insight into how animals make escape decisions in complex, real-world environments.

## Introduction

Wild animals constantly face threats from predators. Rapid threat detection and response are crucial for survival [1]. Prey must rapidly assess their surroundings and initiate escape when necessary in order to reach safety in time [2]. While these assessments are challenging, prey are remarkably successful at doing so, a phenomenon reflected in the low capture success rates of predators reported in aquatic systems [3–5].

However, escape decisions come with a trade-off. Unnecessary responses waste energy and interrupt foraging opportunities [6]. Failing to respond can be fatal, but fleeing too often wastes a lot of valuable energy. Therefore, successful decision-making depends on reliably distinguishing between threatening and non-threatening situations, as well as adjusting one’s behavior based on information about the surroundings [2].

One of the ecosystems where these decisions are frequently made is the coral reef, where encounters between predators and prey are common [7]. Due to the complex visual environment of the surroundings, filled with conspecifics, non-predators, and a diverse range of predators, escape decisions are multifaceted and challenging. Additionally, the predators inhabiting the coral reef employ diverse predation strategies, including pursuit, ambush, and stalking [8, 9]. While these strategies are vastly different in nature, one thing they have in common is the fast approach of a predator at some point during the attack. For prey to survive, they must be able to detect and quickly respond to these attacks while suppressing escape responses in non-threatening situations.

Fish have several strategies by which they can perceive attacks by predators, including rapid pressure changes [10], sounds [11], and visual cues [12]. Among these, visual cues are particularly important for triggering escape responses in many fish species [12, 13]. Laboratory studies in model fish species have identified specific neural circuits in the brain that respond exclusively to fast visual looming stimuli (an expanding image representing an approaching object) [14, 15]. Looming stimuli have also been shown to trigger escape responses in various animal taxa, including amphibians, birds, insects, and humans, suggesting that while the underlying neural mechanisms may differ, the behavioral response to looming is widespread [15, 16].

The speed of the approaching object is a critical cue influencing the activation of neural circuits, with fast-approaching objects eliciting rapid, reflexive responses through the Mauthner cell pathway [17, 18]. In fish, the Mauthner cells generate a fast, unilateral motor response by activating muscles on one side of the body, causing a sharp bend in the body (referred to as a C-start), after which the fish can swim away in the opposite direction of the threat [19].

In addition to high sensitivity to fast-approaching objects, this pathway has the ability to integrate information about the environment outside of the stimulus, such as the presence of a refuge or other prey animals [20]. Slow-approaching objects are less likely to activate the Mauthner cell pathway, allowing the fish to integrate more information about the object before deciding on the escape response [21–23].

Laboratory studies clearly show that the speed of a looming object strongly influences the probability of a C-start, with fast-approaching objects more likely to elicit rapid escape responses [17, 18]. However, it remains unclear whether these neural circuits can reliably distinguish between visual cues associated with non-threatening movements, such as the cruising speed of predators, and truly threatening motion, such as attack speeds. This distinction is particularly important on coral reefs, where predators are frequently within the visual detection range of prey but rarely launch attacks [7]. In such environments, responding to every approach would generate excessive false alarms, so prey must finely tune their sensitivity to the speed of potential threats.

While the features of an approaching stimulus are central to triggering escape responses, ecological and social contexts also influence these behaviors [20, 24]. Social contexts, such as group formation, function as anti-predatory strategies and often shape escape responses [25–27]. The presence of other prey can reduce the probability of escape responses to external stimuli, even in species that typically forage alone, as seen in some reef-dwelling fish [24]. This suggests that sensitivity to looming objects is adjusted based on local risk, a principle that may extend beyond social context alone. Many coral reef fish show strong site fidelity and depend on coral structures for shelter, which influences their risk of predation [28, 29]. Individuals located farther from the shelter face a greater risk, as returning takes longer. Yet the interaction between ecological and social factors, threat-related cues, and species differences remains unclear, particularly under natural conditions.

This research aims to unravel the behavioral rules individuals use when making escape decisions in complex environments by exposing groups of site-associated coral reef fishes to simulated predator attacks of different speeds and testing the importance of social and environmental contexts. Additionally, the swimming speeds of local predators during normal cruising and active attacks are analyzed to test whether prey responses are tuned to real predation events. By combining high-resolution behavioral tracking with naturalistic experiments, this study reveals how prey integrate sensory input with environmental context to make rapid, adaptive decisions in complex, predator-rich ecosystems.

## Materials and methods

### Data collection and video processing

#### Experimental setup

The experiments were conducted on shallow reef flats at Cas Abou and Kokomo Beach along the west coast of Curaçao, in the Caribbean Sea (Fig 1A). A total of 12 trial sessions were conducted at 5 sites around the reef, all located at approximately 3 meters in depth. Each trial session took place on an isolated coral patch that serves as a refuge for various fish species. Since most fish remained close to the coral, we could reliably track individuals during each stimulus presentation.

**Fig 1.**
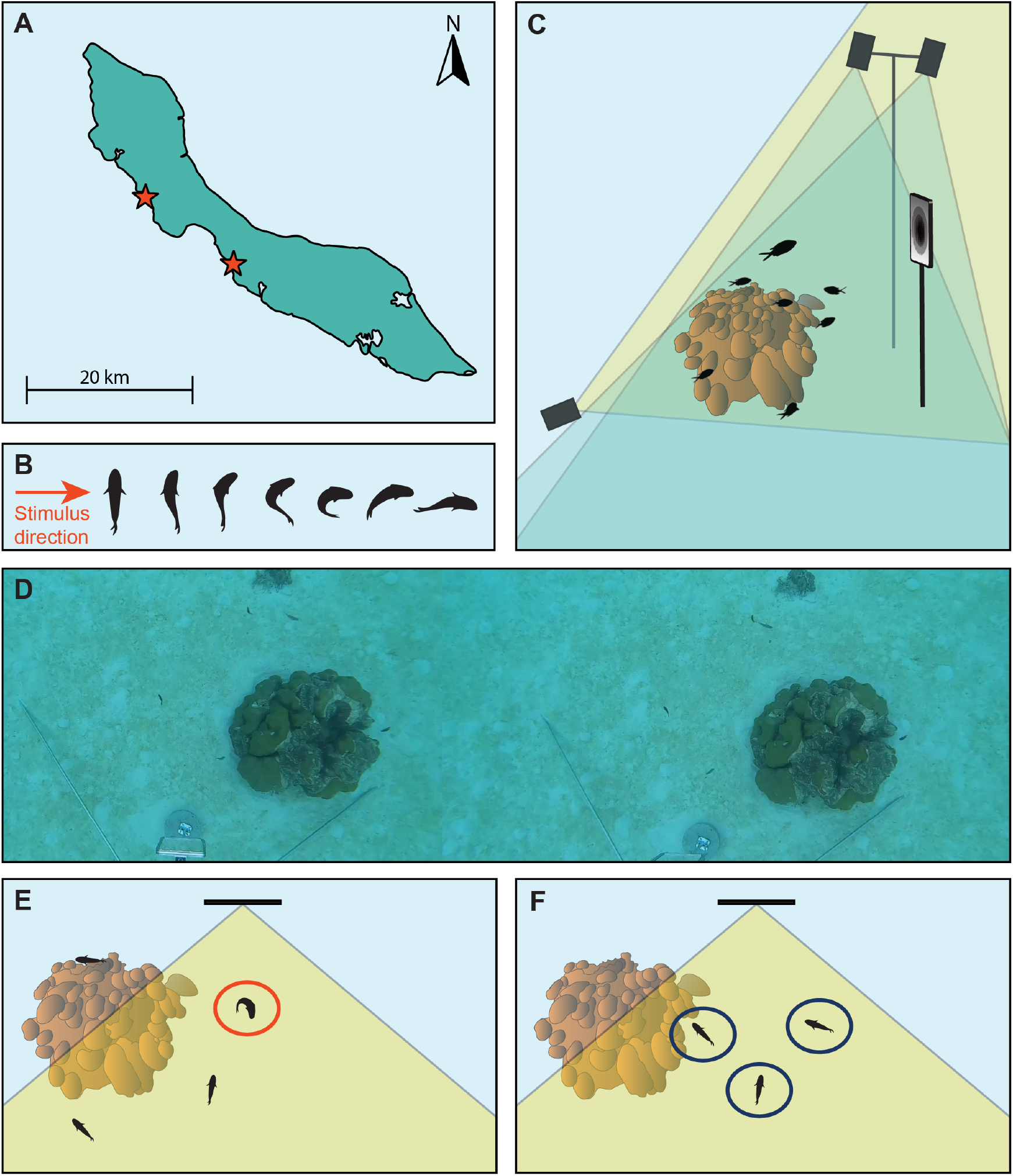
Overview of experimental setup. A: Locations of setup in Curaçao. Red stars mark the experimental sites at Cas Abou and Kokomo Beach. B: Top view of body postures during escape response (referred to as a C-start [19]). C: Experimental setup around an isolated coral patch, with two top-view cameras (blue line of sight) and one side-view camera (yellow line of sight). D: Top view from the left and right top-view cameras. E: First-responding fish (FR; red circle) in response videos are compared with F: Non-responding fish that see the stimulus (NRL-NR; blue circles) in the non-response videos.

The two dominant species, Brown Chromis (*Chromis multilineata*) and Bicolor Damselfish (*Stegastes partitus*), typically settle at these coral refuges during their juvenile stage and remain there until adulthood [30]. In addition to these species, the coral patches are frequently visited by other reef-dwelling species that belong to various families of coral reef fish.

At the experimental sites, we set up an iPad (iPad Pro 2022, 12.9 inches) to display a looming stimulus every 90 seconds, for a total duration of one hour per trial session. Three cameras (GoPro HERO9, 1080p, 240fps) were positioned around the coral patch: two facing downward and one arranged for a side view facing the stimulus (Fig 1C). The looming stimulus consisted of an expanding black dot on a white background, simulating an approaching object (a predator) in the region in front of the iPad screen. The expansion rate of the dot was calibrated to match the optical expansion produced by a predator approaching at one of nine different speeds (0.5, 1.0, 2.0, 3.0, 4.0, 5.0, 6.0, 7.0, or 8.0 m s^*−*1^), and the order of the loom stimulus speeds was randomized across presentations within each trial session. For a more detailed description of the looming stimulus, see S1 Appendix (Looming stimulus).

#### Video analysis and 3D reconstruction

The video data from the three cameras were synchronized, and all recordings were manually screened for the presence of escape responses. An escape response was defined by a body bend (a C-start; Fig. 1B) followed by a rapid increase in speed [19]. For each fish exhibiting an escape response, the frame corresponding to the C-start was recorded (Fig. 1D).

To reconstruct the 3D positions of the fish and their environment, 2D positions from the two downward-facing cameras were manually annotated using the open-source platform CVAT [31]. These 2D positions were triangulated into 3D coordinates based on stereo correspondence between paired cameras using calculated stereo parameters [32]. To obtain these parameters, the cameras were calibrated before each trial session by presenting a calibration checkerboard at various orientations and positions within the cameras’ field of view. Stereo calibration parameters were estimated using the Stereo Camera Calibrator app in MATLAB [33]. For a detailed description of the environmental reconstruction, see S2 Appendix (Coral reconstruction).

Each fish was assigned a unique ID, and its species and response frame (if applicable) were recorded. Fish-specific variables, such as distance to coral, were calculated using the reconstructed 3D positions of the fish and the environment. Some fish were excluded from analysis because they were not visible in one of the stereo cameras or left the field of view, which prevented the calculation of their 3D positions or speeds. Videos were excluded when more than 33% of the tracked fish were unusable. After filtering, the dataset comprised 67 videos with at least one responding fish and 180 videos with no observed responses.

#### Measuring natural attack speeds

To quantify the attack speeds of a natural predator of the two focal prey species, stereo cameras (GoPro HERO9, 1080p, 120fps) were deployed in the same coral reef as the experimental setup. From these recordings, 61 attacks by Bar Jacks (*Caranx ruber*) were isolated and analyzed. Each attack was tracked from the moment the Bar Jack entered the field of view or initiated an attack until the prey, typically a Brown Chromis (*Chromis multilineata*), either evaded capture or was caught.

3D tracks of the Bar Jack and Chromis were manually annotated by tracking head positions. Attack speed data were computed by calculating the magnitude of the change in position per frame and dividing it by the frame rate. The 90th percentile of the predator’s instantaneous speed was used as a proxy for maximum attack speed. This measure captures the fast-approach phase of the attack, which is most relevant to prey escape decisions, while reducing sensitivity to tracking noise associated with using absolute maxima.

### Calculation of variables

To analyze how different informational contexts influenced escape decisions, we calculated a set of explanatory variables (see Table S1 for an overview). These variables describe both experimental manipulations and natural variation in the context of the fish. Data processing was conducted in Python (version 3.13.6), and model fitting and statistical analyses were performed in R (version 4.5.1).

The programmed speed of the looming stimulus (LSp) was determined by the experimental design and serves as the primary predictor variable. We chose to use the programmed speed rather than the calculated perceived speed or size. Perceived speed changes continuously as the stimulus approaches and requires a specific time point at which to be evaluated. For responding fish, this reference point is unambiguous (the frame of the C-start). For non-responding fish, no equivalent reference frame exists, making it impossible to assign a comparable perceived speed value.

The order of stimulus presentations within a trial (trial event number; TEv) was included to test for habituation, defined as a reduction in responsiveness to repeated stimuli. While habituation (a decrease in response to repeated stimuli) is commonly observed in laboratory animals [34], it tends to be slower and more context-dependent in wild animals [35].

Several variables captured how well each fish could perceive the stimulus. The distance to the stimulus (DSt) was measured as the three-dimensional distance between the fish’s head and the center of the iPad screen. Because the apparent size of an object scales with the inverse square of distance, fish farther from the iPad experienced a much smaller visual stimulus than those nearby [36]. The viewing angle described the angle at which the fish viewed the iPad screen. A viewing angle of 0^*°*^ corresponds to a fish directly in front of the iPad screen, with the angle increasing as the fish moves more to the side. At larger angles, the screen appeared dimmer, and it became reflective at a large enough angle (≳ 50^*°*^) due to a Snell’s window effect [37]. The orientation angle (OA) describes the fish’s body alignment relative to the stimulus, ranging from 0^*°*^ when facing the iPad directly to 180° when facing directly away. This variable was included to account for the possible effect of predator approaching angle on escape probabilities.

Species identity (Sp) was manually identified from the video footage and included to assess interspecific differences in escape behavior. Distance to coral (DCo) was measured as the shortest three-dimensional distance to the nearest coral patch, which serves as a potential refuge. Neighbor Proximity (NPro) was calculated as the sum of the inverse squared distances to neighboring fish, giving nearby individuals greater influence than those further away. This weighting reflects that both the apparent size of a neighbor’s image on the retina and its potential behavioral influence decrease with the square of distance [13, 36, 38].

### Dataset: classification of individuals

The experiment allowed the classification of fish into distinct groups based on the information available to them. To ensure that the fish responded only to the stimulus, we compared the first responders in the response videos (FR, Fig 1E) with the unresponsive fish that saw the stimulus in the non-response videos (NRL-NR, Fig 1F). Both groups consisted of fish with non-responding neighbors, ensuring their response behavior was not influenced by the responses of others. For a complete description of the classification, see S3 Appendix (Classification of individuals).

Fish in videos with a response were categorized based on their exposure to the stimulus and the frame of their response. First responders (FR, n = 95) included all fish that initiated a response and any secondary responders that responded within five frames (20.83 *ms*) after the first responder. This time frame accounts for the sensory-motor delay required for visual information processing [39] and was determined as the minimum interval between a first responder and a secondary responder that could not directly see the stimulus in a video. For videos without any escape responses, individuals within a viewing angle of 55° were classified as fish that saw the stimulus but did not respond (NRL-NR, *n* = 1255). This threshold corresponds to the approximate limit of visibility caused by the Snell’s window effect, beyond which the iPad screen becomes reflective.

### Model selection

#### Random effect

The dataset comprises data from 12 trial sessions conducted at five different reef sites. The number of data points varied across trial sessions, sites, and site repetitions (see Fig S1, panels A and C). In addition, the different trial sessions varied strongly in the species composition (Fig S1, panel B), as well as the neighbor proximity of the fish (Fig S1, panel D). To account for the unbalanced dataset, the specific trial session ID was used as a random effect in the model selection and fitting process (see S5 Appendix for statistical justification).

#### Linear effects of single variables

All analyses were performed using generalized linear mixed models (GLMMs) with a binomial error structure, logit link function, and a random effect for the trial session ID. Continuous predictors were scaled to have a mean of zero and a standard deviation of one to standardize effect sizes and improve model convergence.

To assess how each predictor influenced response probability, we first fitted single-predictor models with a random intercept for trial session. We then fit a multivariable model including all predictors simultaneously to account for their joint effects.

#### Non-linear effects of single variables

To account for potential threshold effects or other non-linear relationships between predictors and response probability, each continuous variable was tested in multiple functional forms by replacing it in the multivariable model with alternative transformations. Each predictor was tested in four forms: linear, logarithmic, and natural splines with two and three degrees of freedom, while all other predictors were initially held in their linear form. Model fits for these alternatives were compared using the Bayesian Information Criterion (BIC) to identify the best-fitting transformation for each variable. To evaluate the influence of non-linear transformations in the context of other non-linear variables, each variable was tested in a multivariable model where the other variables were in their optimal functional form.

#### Interaction effects

To identify potential interactions between predictors, a list of ecologically relevant interactions was systematically tested using a multivariable model that incorporated the previously identified non-linear transformations of individual predictors. Each interaction term was added individually to the base model and compared to it using a likelihood ratio test (LRT) to evaluate whether the inclusion of the interaction term significantly improved the model fit. To account for multiple testing and control the false discovery rate, the raw p-values were adjusted using the Benjamini-Hochberg procedure [40].

To limit the number of interaction terms and focus on biologically meaningful combinations, we considered only two-way interactions among four ecologically relevant variables: looming stimulus speed (LSp), distance to the coral refuge (DCo), species identity (Sp), and neighbor proximity of neighboring fish (NPro). These variables were chosen based on a priori expectations that their effects on escape responses might be modulated by one another, reflecting trade-offs between environmental risk, social context, and species-specific behavior.

#### Species differences

To evaluate whether species differences in escape probability could be attributed to other predictors, the effect of additional predictors on the species effect was tested. The species-only univariable model was compared to the same model extended with each individual predictor. For each model, the Akaike Information Criterion (AIC) and species coefficient estimates were compared. A reduction in the species coefficient, or a loss of significance, was interpreted as evidence that the added predictor accounted for part of the species effect. AIC was used here rather than BIC, as the goal was to assess changes in the species coefficient rather than to select between competing model structures.

## Results & Discussion

Our results show that escape responses were primarily dominated by the speed of the stimulus, and further influenced by spatial positioning and species identity. Contrary to expectations, social context had no measurable effect. Together, these results suggest that escape decisions in reef fish are highly sensitive to information about the threat itself, while ecological factors play secondary roles in influencing these decisions.

### Threat characteristics dominate escape decisions

#### Stimulus speed is the primary driver of escape responses

Across all analyses, stimulus speed (LSp) emerged as the strongest predictor of escape responses. In the linear univariable model, higher stimulus speeds significantly increased response probability (*β* = 1.25 *±* 0.15, *p* = 7.0 *×* 10^*−*18^; Table S2), and the effect remained highly significant in the multivariable model (*β* = 1.37 *±* 0.17, *p* = 3.2 *×* 10^*−*16^; Table S3). Among all univariable models, it explained the largest proportion of variance 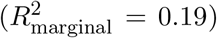. Response probability increased non-linearly with stimulus speed, showing a sharp rise above approximately *≈* 2.0 m s^*−*1^ (Fig. 2A), which was supported by model comparisons indicating that non-linear forms provided a better fit than the linear form (Table S4).

**Fig 2.**
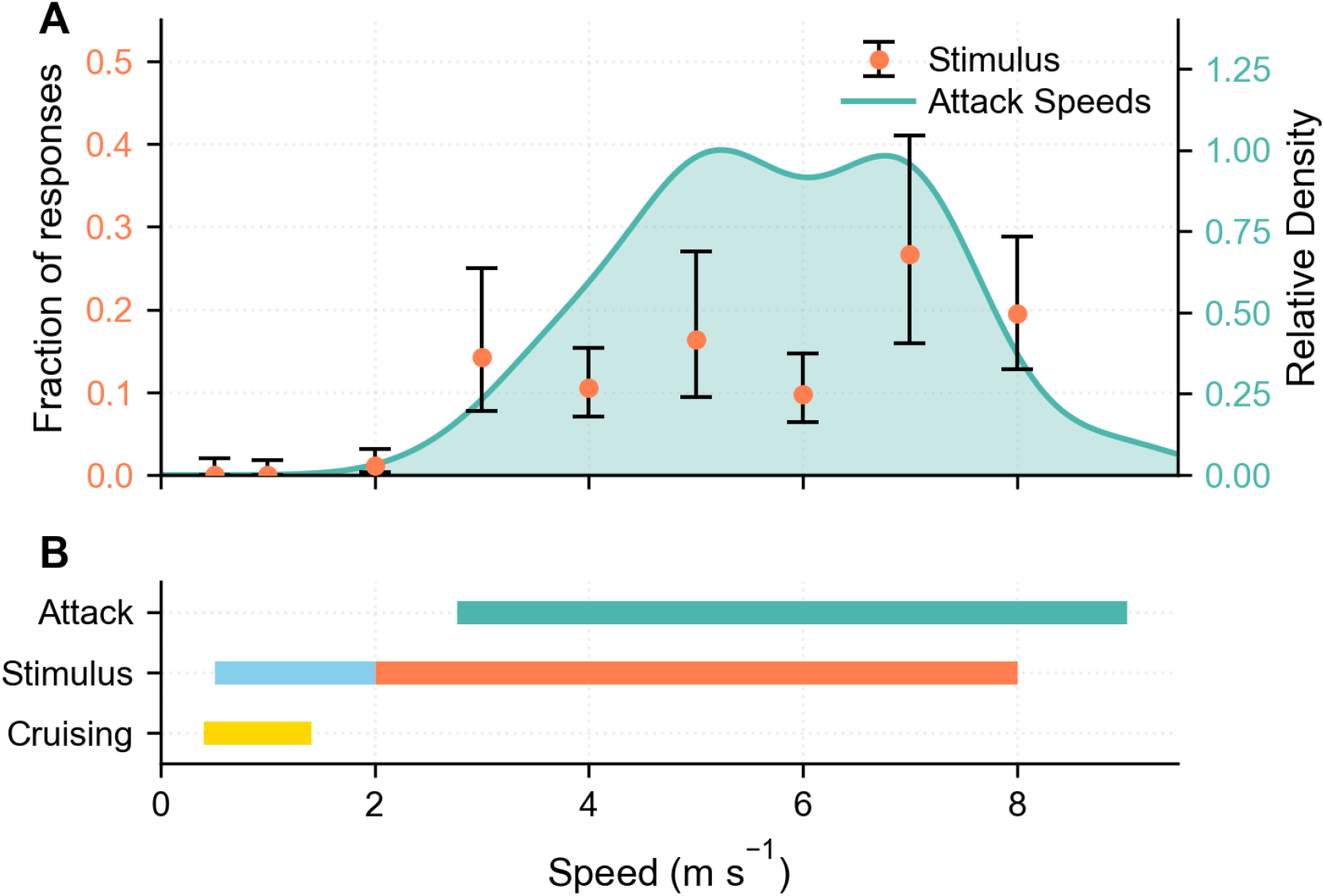
Responses per stimulus speed and comparison with natural swimming speeds of predators. A: The fraction of stimulus presentations at each speed that resulted in an escape response (orange points *±*95% CI). Overlaid is a kernel density estimate of the 90th-percentile attack speeds from 61 recorded Bar Jack attacks (teal), representing the distribution of natural predator attack speeds. B: Speed ranges of natural attack speeds (teal) overlap with stimulus speeds that triggered escape responses (orange). Stimulus speed at lower speed ranges (light blue) did not trigger any responses, and overlapped with natural cruising speeds of predators (yellow).

This threshold-like pattern indicates that fish were largely unresponsive to slow-moving stimuli but responded reliably once the stimulus exceeded a certain speed. Such non-linear sensitivity is consistent with the activation properties of the Mauthner-cell escape circuit, which is tuned to rapid visual expansion [21–23]. This tuning may allow fish to avoid unnecessary escapes in non-threatening situations while maintaining high responsiveness to fast, potentially dangerous movements.

#### Speeds that trigger responses align with natural predator attack speeds

Escape responses were only observed when stimulus speed exceeded 2.0 m s^*−*1^, with the majority occurring at speeds of 3.0 m s^*−*1^ or higher (Fig. 2A). This behavioral threshold closely aligns with the attack speeds of Bar Jacks (*Caranx ruber*), which ranged from 2.7–9.0 m s^*−*1^ (90th percentile; Fig. 2B). Routine cruising speeds for *C. ruber* have not been reported, but the routine swimming speed of another carangid (e.g., Green Jack (*Caranx caballus*)) indicates routine speeds on the order of *≈* 1 m s^*−*1^ or less for similarly sized fishes. These values are consistent with broader comparative studies in reef fishes, showing that attack movements are several times faster than routine swimming [41].

The close correspondence between response thresholds and natural predator attack speeds indicates that fish selectively respond to the motion characteristics of attacks rather than ordinary cruising movements. This supports the idea that animals use the speed of approaching objects as a key cue for threat discrimination [15, 16, 24]. Avoiding responses to slower stimuli may help reduce false alarms in ambiguous situations, as slower movements are more likely to represent non-threatening situations.

#### Spatial positioning influences visual access to the stimulus

Variables that were specific to the experimental setup (specifically the distance to the stimulus (DSt) and the angle at which fish viewed the iPad (VA)) had a significant role in determining whether a fish initiated an escape response. Fish closer to the screen and viewing it from a smaller angle had a higher probability of responding.

These effects likely reflect the perceptual geometry of the setup. Because the looming stimulus was presented on a flat screen, it was most clearly visible when viewed from close range and at a smaller angle. At greater distances or wider angles, the screen became more reflective and the stimulus less distinct, making the threat harder to detect. This pattern is consistent with laboratory findings showing that response probability declines under reduced visual contrast and suboptimal viewing geometry [20, 37].

For a detailed description of the effects, see S6 Appendix (Effect of experimental variables).

### Species employ different antipredator strategies linked to spatial positioning

Species identity had a significant influence on the likelihood of escape responses. In the univariable model, Bicolor Damselfish (*Stegastes partitus*) had a substantially lower response probability than Brown Chromis (*Chromis multilineata*) (*β* = *−*1.66 *±* 0.37, *p <* 0.001; Table S2). This effect remained significant in the multivariable model (*β* = *−*1.10 *±* 0.48, *p* = 0.02; Table S3).

However, the two species differed in their distance to the coral refuge. Brown Chromis tended to be farther from the coral (Fig 3B), while Bicolor Damselfish were typically located closer to shelter (Fig 3C). This difference was significant (Wilcoxon rank-sum test, *W* = 351,766, *p <* 0.001), confirming the correlation between species identity and distance to the coral. Distance to coral (DCo) itself had a significant positive effect on response probability in the univariable analysis (*β* = 0.64 *±* 0.12, *p <* 0.001; Table S2), indicating that fish farther from shelter were more likely to flee (Fig 3A).

**Fig 3.**
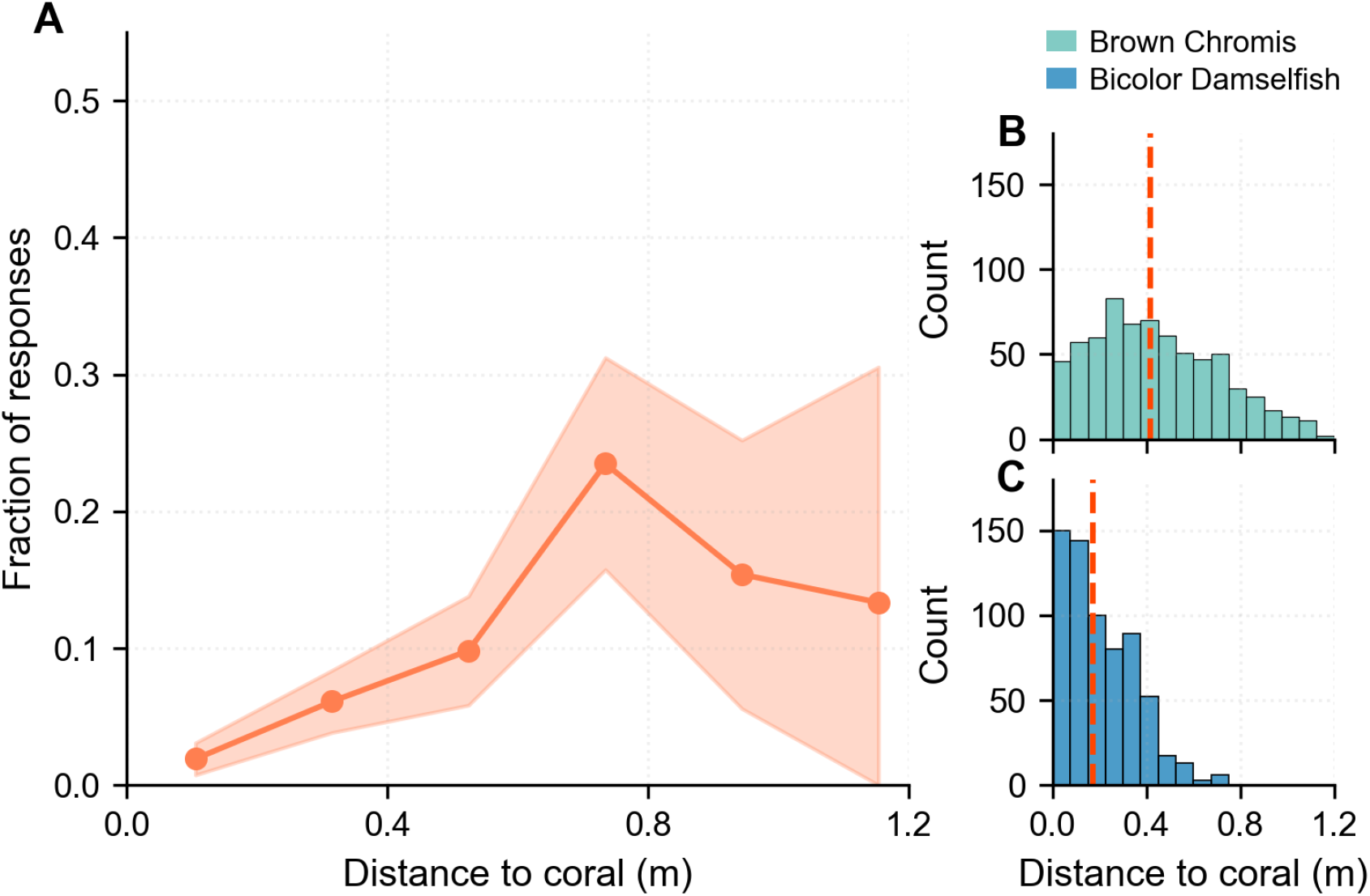
Species differences in escape responses are linked to distance from coral refuge. A: Fraction of fish responding across distance-to-coral bins, with shaded areas showing 95% CIs. Response probability increased with distance to the coral shelter, peaking at intermediate distances. B: Distribution of distances to coral for Brown Chromis. Red dashed line indicates median distance. C: Distribution of distances to coral for Bicolor Damselfish. Red dashed line indicates median distance.

When both predictors were included in the same linear multivariable analysis, the effect of distance to the coral lost its statistical significance (*β* = 0.11*±*0.17, *p* = 0.54; Table S3), although a log-transformed version improved model fit (ΔBIC = *−*4.92; Table S4). In the final non-linear version, the effect remained non-significant (*β* = 0.29 *±* 0.22, *p* = 0.172; Table S5). Including the log-transformed form in a species-only model substantially reduced the species coefficient (from *β* = *−*1.66 to *β* = *−*1.03) and improved model fit (ΔAIC = *−*18.4), but the species effect remained statistically significant (*p* = 0.01; Table S6).

These results suggest that the two species use different antipredator strategies. Brown Chromis appear to employ a high-risk, high-vigilance strategy, where they stay further from the shelter but compensate with high responsiveness. Bicolor Damselfish follow a low-risk, shelter-dependent strategy, remaining closer to the refuge and relying less on flight responses.

The persistence of species-specific differences after accounting for distance to coral indicates that these behavioral patterns are shaped by intrinsic traits rather than by refuge availability alone. Bicolor Damselfish are known for their territorial and aggressive behavior [42], and may prioritize defending their coral patch over fleeing. Brown Chromis, by contrast, are more mobile and less tied to specific shelter sites, allowing them to forage farther from shelter while relying more heavily on rapid escape when threatened. Their greater distance from shelter likely reflects foraging opportunities in the water column, where planktonic prey are most abundant [28, 29].

### Social context had no significant effect on response

Contrary to expectations, the proximity of neighboring fish did not have a significant influence on the escape probability. In the univariable analysis (Table S2), no effect was detected 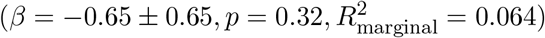. In the multivariable model (Table S3), the effect remained non-significant (*β* = *−*0.62 *±* 0.74, *p* = 0.40). None of the tested interaction terms of neighbor proximity (involving distance to coral, species, and stimulus speed) improved the model fit (all adjusted *p ≥* 0.324, Table S7).

We also tested two alternative measures for social context (the total angular area of neighbors in the focal fish’s visual field (NAS) and in the visual hemisphere containing the stimulus (N). Neither measure showed a significant effect in the multivariable analysis (see S7 Appendix), supporting the conclusion that social context did not affect escape responses.

These findings are surprising, as higher neighbor proximity has been shown to suppress responses in other studies, attributed to the reduced per-capita risk through dilution effects [24–27]. A possible explanation for this lack of social effects is that the fish in this study were much smaller than those in previous research [24]. For small fish, a large number of neighbors is needed to create a meaningful dilution effect. From direct observation during fieldwork, we noted that aggregations of these species can reach substantially larger sizes than the loosely aggregated groups observed during this study. As the fish in this study were loosely aggregated, the local densities may have been insufficient to give rise to the dilution effect seen in denser groups.

### Other variables did not affect the response

The other variables tested in this analysis had no significant effect on the response probability. The exact orientation of the fish relative to the stimulus (OA) did not have a significant effect on the response probability (Fig S4A), which is consistent with the fact that fish have a visual field of almost 360° [43].

The order in which the stimulus was presented (Trial event; TEv) showed statistically significant spline terms in the final non-linear model (Table S5), but the overall improvement in model fit compared to a linear form was negligible (ΔBIC = *−*1.34; Table S4). In the linear multivariable model, the trial event number was not significant (*p >* 0.05; Table S3), and the trend in response probability across successive trials did not show a consistent decline (Fig S4B). Together, these results suggest that habituation did not occur within the time frame of our study, consistent with the idea that habituation is limited under natural field conditions where threats are unpredictable and variable [35]. The statistically significant spline terms likely reflect minor fluctuations across the stimulus showings rather than a systematic change in responsiveness.

Finally, none of the prespecified two-way interactions (Table S7) significantly improved model fit after controlling for the false discovery rate (Benjamini-Hochberg). The interaction between stimulus speed (LSp) and distance to the coral (DCo) lost its significance after the correction (*p*_adjusted_ = 0.165), and all others were non-significant in all analyses *p ≥* 0.324).

## Conclusion

This study demonstrates that escape decisions in reef fish are primarily driven by visual information about the threat itself. Responses only occurred when stimulus speeds matched those of natural predator attacks, suggesting that the looming-sensitive visual circuit is highly tuned to distinguish threatening motion from non-threatening motion. This close correspondence indicates that effective threat discrimination can emerge from relatively simple perceptual processing rather than complex cognitive evaluation.

The effect of the stimulus speed was consistent across species. However, species-specific strategies further shaped escape behavior. Bicolor Damselfish, which are territorial and shelter-dependent, stayed closer to the coral and had a lower response probability. Brown Chromis, which are more mobile and less site-bound, employed a high-risk, high-vigilance strategy, foraging further from shelter but compensating with higher responsiveness.

Surprisingly, the social context had no measurable effect on escape decisions, contrasting with previous research findings. This suggests that for small, loosely aggregated reef fish, dilution effects may not emerge if group sizes are too small to reduce predation risk meaningfully.

These findings highlight that escape behavior in reef fish relies on a conserved perceptual mechanism for detecting threats, which is finely tuned to distinguish true danger from non-threatening motion. Species-specific strategies further determine how individuals respond to this information. These results demonstrate that reef fish decision-making relies on conserved looming-detection mechanisms finely tuned to predator attack speeds.

## Acknowledgments

We thank Lars Koopmans for his contributions to data collection and processing. Additionally, we thank the staff at the Department of Theoretical and Computational Ecology for providing feedback during our meetings, in particular, Prof. Dr. André de Roos. We are grateful to the authorities of Curaçao for granting research permissions (permit number: #2022/21467).

## Author contributions

Conceptualization: SN, BM. Data curation: BM, SN, LF. Formal analysis: SN, LF. Funding acquisition: BM. Investigation: BM. Methodology: BM, SN. Project administration: SN, BM. Resources: BM. Software: SN, LF. Supervision: BM. Validation: SN. Visualization: SN. Writing - Original draft preparation: SN. Writing - Review & Editing: SN, BM.

## Competing interests

The authors declare that they have no competing interests.

## Data availability

All data and analysis code used in this study are publicly available on GitHub at: https://github.com/saralneven/InitialResponders. A permanent snapshot of the repository is archived at Zenodo: https://doi.org/10.5281/zenodo.18989233.

## Ethics statement

All fieldwork was conducted under the Curaçaoan Government’s Permit #2022/21467 to CARMABI. The reef sites are publicly accessible. Because no animals were handled, this research was determined not to require a license from the Head Animal Welfare Body of the University of Amsterdam, as set out in the Dutch Experiments on Animals Act.

## Funding statement

This research was supported by the Netherlands Organization for Scientific Research (NWO grant VI.Vidi.203.085). The funders had no role in study design, data collection and analysis, decision to publish, or preparation of the manuscript.

## Supporting information

**Fig S1.**
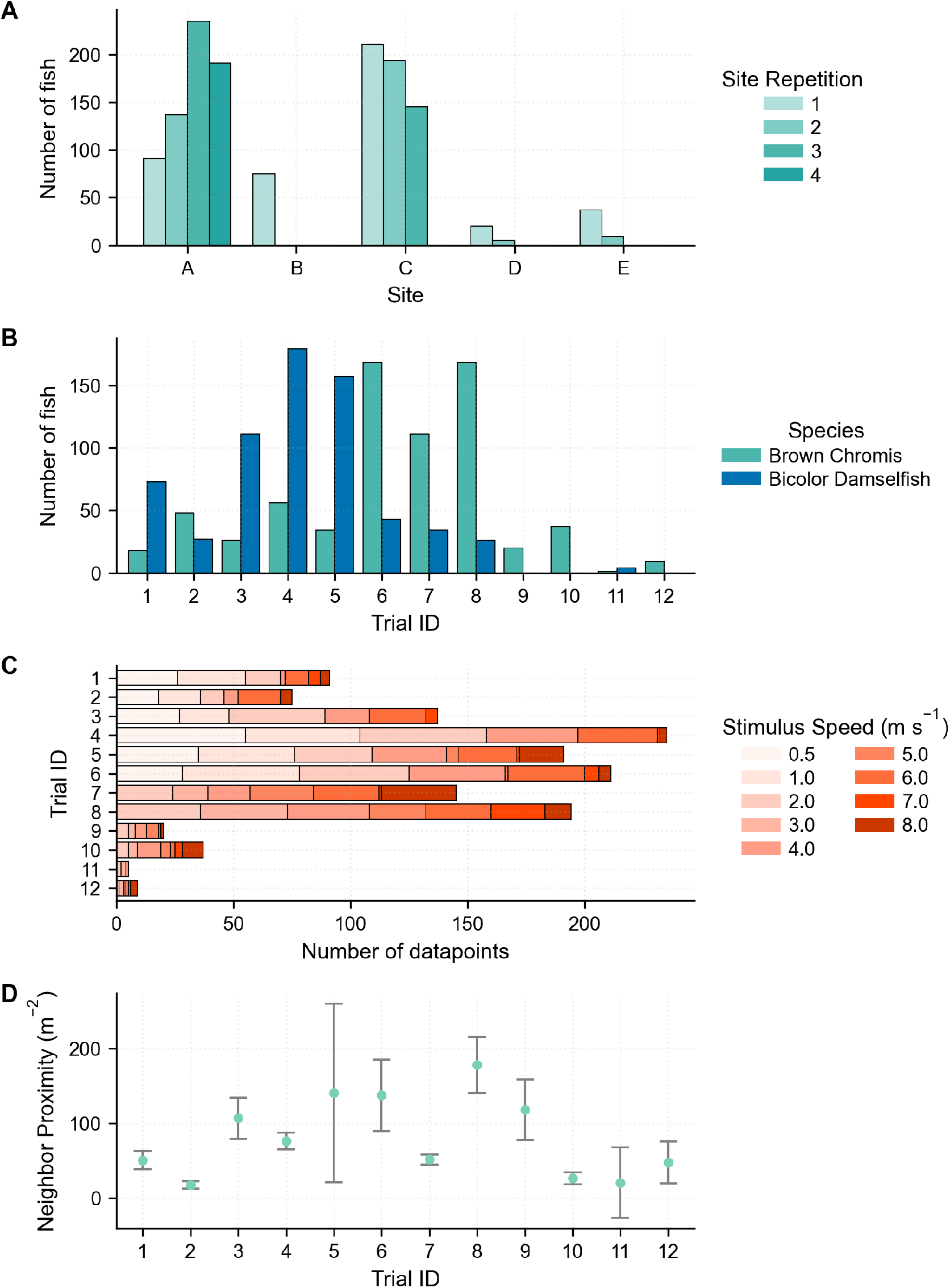
Variation between trial sessions. A: Number of fish per site repetition. B: Number of fish per species across trial sessions. C: Number of datapoints per trial session. D: Neighbor proximity across trial sessions.

**Fig S2.**
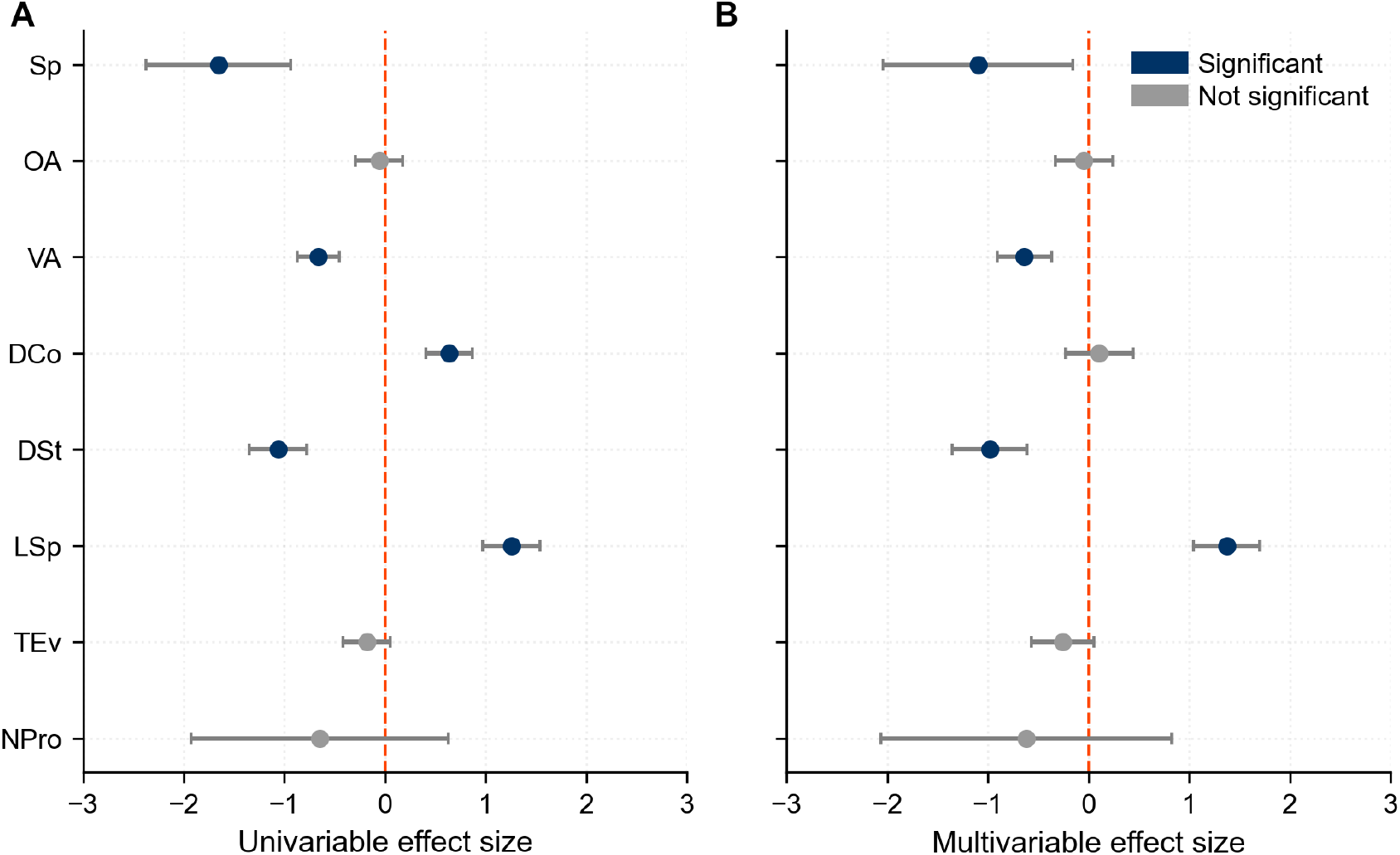
Effect sizes and significance in linear models. Effect size estimates (*β*) and 95% CIs from (A) univariable models and (B) the multivariable model. Significant predictors are shown in dark blue, non-significant in gray.

**Fig S3.**
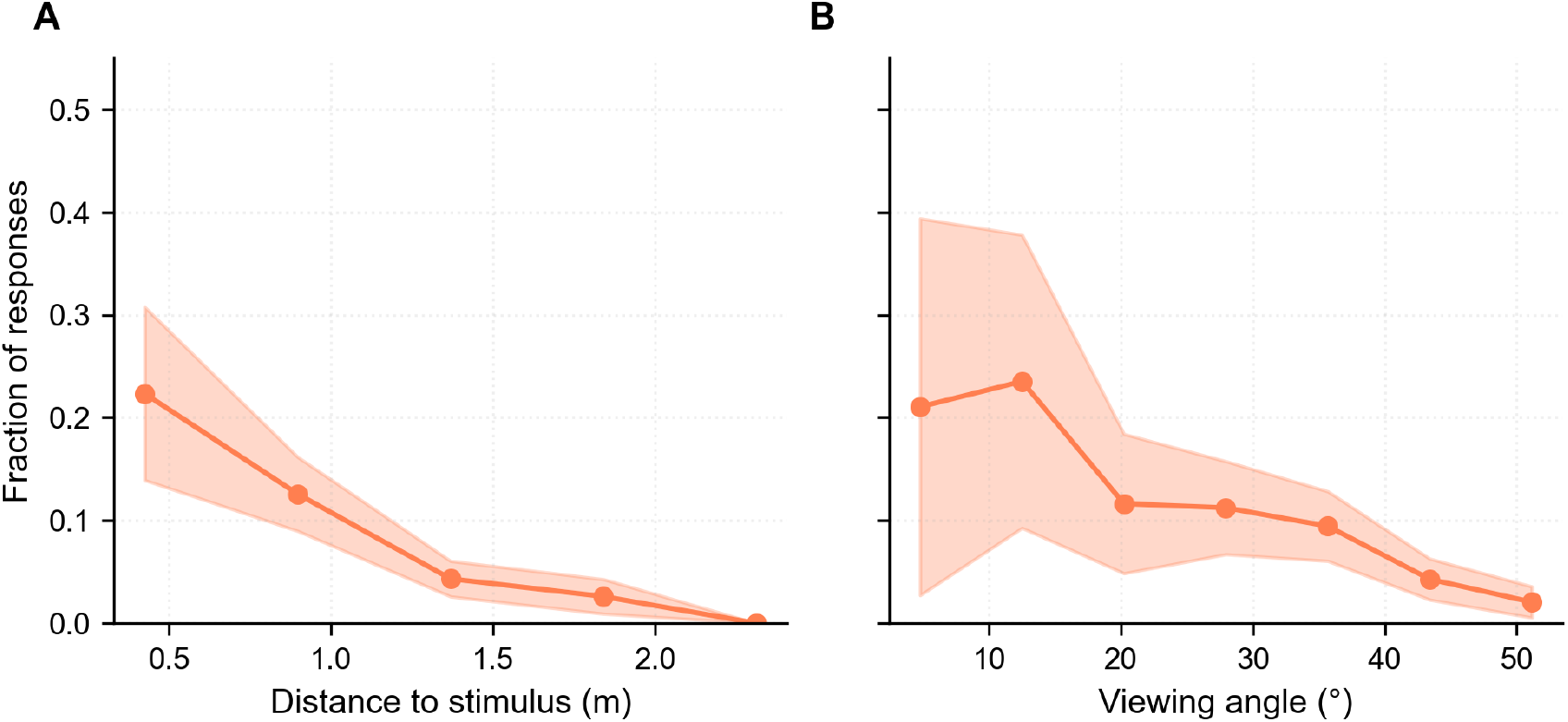
Escape response probabilities decline with distance to the stimulus and viewing angle. (A) Fraction of responses across distance to stimulus bins. (B) Fraction of responses across viewing angle bins. Shaded areas represent 95% CIs.

**Fig S4.**
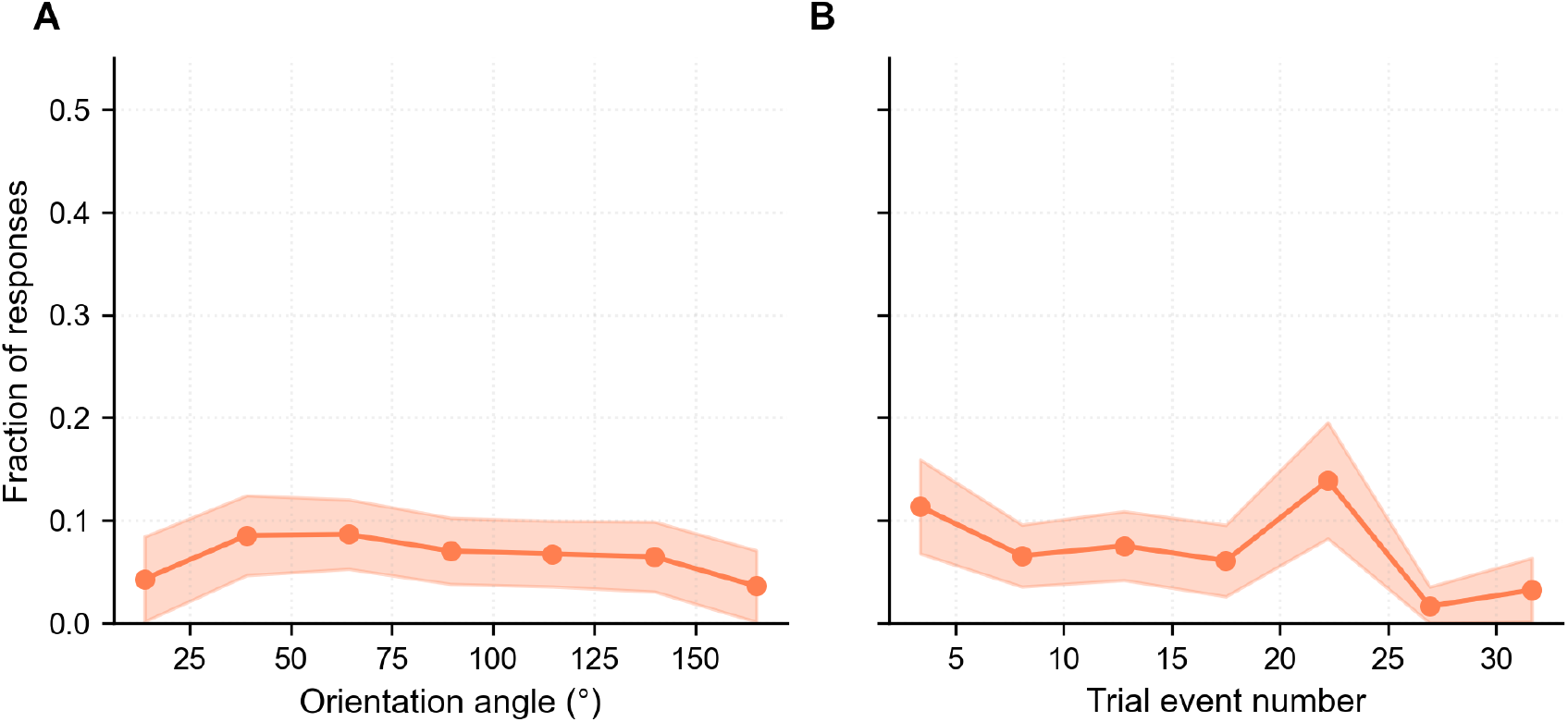
Effects of orientation angle and trial event number. (A) Fraction of responses across orientation angle bins. (B) Fraction of responses across trial event number. Although spline terms were statistically detectable, the effect was weak and inconsistent.

## S1 Appendix Looming Stimulus

We generated looming stimuli to simulate a predator approach using a black circle that expanded on a white background. When viewed from a distance of 1 meter in front of the iPad screen (*d*_screen_), the looming image corresponded to a predator with a radius of 4 cm (*r*_predator_ = 0.04 m). Based on typical depth–length relationships for reef fishes [44], this radius corresponds to a predator approximately 30 cm in total length, which reflects a typical predator size for the 6–10 cm Brown Chromis and Bicolor damselfish in our study. We simulated attacks with varying predator speeds (*v* = 0.5–8.0 m s^*−*1^) from a starting distance (*D*_0_ = 10 m). At the initiation of each simulated attack, the radius of the looming circle (*R*_loom_, meters) at time *t* was calculated as:

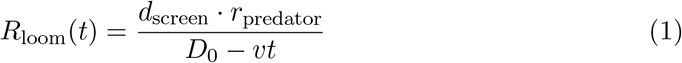

where *t* is time (s). To generate the image of the black circle, the calculated radius was converted to pixels using the pixel density of the iPad (10,394 px m^*−*1^). Videos of the looming stimuli were generated and played back at 120 frames per second.

## S2 Appendix Coral reconstruction

Annotation of the coral was done using the same annotation software as for the fish (CVAT.ai Corporation, [31]). For every trial, multiple points on the coral were annotated and converted to 3D points. Three-dimensional reconstructions of coral patches were generated from annotated point clouds. The point sets were processed using a Delaunay-based triangulation, which connected neighboring points to form a mesh of triangular faces. Duplicate faces were removed to obtain a clean surface, resulting in a triangulated structure that accurately captured the overall geometry of each coral patch. This triangulated surface provided a compact, geometry-preserving representation of each coral patch suitable for spatial analyses.

## S3 Appendix Dataset Classification of individuals

### Response videos

Fish in videos with a response were categorized based on their exposure to the stimulus and the frame of their response. First responders (FR, n = 106 before filtering; n = 95 included in this study) included all fish that initiated a response and any secondary responders that responded within five frames (20.83 *ms*) after the first responder. This time frame accounts for the sensory-motor delay required for visual information processing [39] and was determined as the minimum interval between a first responder and a secondary responder that could not directly see the stimulus in a video. Secondary responders were further categorized based on their position relative to the stimulus. Those who viewed the stimulus at an angle smaller than 55° were classified as secondary responders who saw the looming stimulus (SRL, n = 143). Secondary responders outside this 55° angle were classified as secondary responders that could not see the looming stimulus and were, therefore, responding only to social cues from their neighbors (SRNL, n = 141). This distinction reflects the iPad screen’s reflectivity at large angles, which makes it impossible to see the stimulus.

Non-responders were divided into categories based on their visibility of the stimulus as well. Non-responders within the 55° angle were classified as non-responders who saw both the looming stimulus and the escape responses of their neighbors (NRL-R, n = 335). Those outside the 55° angle were classified as non-responders who only observed the escape responses of neighbors (NRNL-R, n = 444).

### Non-response videos

For videos without any escape responses, individuals within the 55° angle were classified as fish that saw the stimulus (NRL-NR, n = 1341 before filtering; n = 1255 included in this study). Individuals outside the 55° angle were classified as fish that had no access to information about the stimulus (NRNL-NR, n = 1434). As no fish responded in the videos, neither type had neighbors responding.

### Final dataset

To ensure that the fish responded only to the stimulus, we compared the first responders in the response videos (FR, Fig 1E) with the unresponsive fish that saw the stimulus in the non-response videos (NRL-NR, Fig 1F). Both groups consisted of fish with non-responding neighbors, ensuring their response behavior was not influenced by the responses of others.

Four FR fish had viewing angles greater than 55°. Two had a viewing angle around 58°, which may reflect minor annotation errors. The other two had angles around 64°, but were positioned very close to the iPad, making the stimulus visible as it increased in size, since the viewing angle was measured from the center of the stimulus. All individuals were excluded from the final analysis.

As the species group ‘Other’ consisted of a variety of species and families with small sample sizes, including Basslets (including groupers and soapfish, *Serranidae*; n = 45), Wrasses (*Labridae*; n = 16), Parrotfish (*Scaridae*; n = 5), Snappers (*Lutjanidae*; n = 6), Doctorfish (*Acanthuridae*; n = 6), Dusky Damselfish (*Stegastes adustus*; n = 5), Sand tilefish (*Malacanthus plumieri* ; n = 2), Flounders (*Bothidae*; n = 2), Angelfish (*Centropyge*; n = 2), Banded Butterflyfish (*Chaetodon striatus*; n = 1), Goatfish (*Mullidae*; n = 1), Boxfish (*Ostraciidae*; n = 1) or Yellowtail Hamlets (*Hypoplectrus chlorurus*; n = 1), the group was excluded from the analysis.

The remaining dataset consisted of two species: Brown Chromis (*Chromis multilineata*; n = 696; FR = 83, NR = 613) and Bicolor Damselfish (*Stegastes partitus*; n = 654; FR = 12, NR = 642), which resulted in 95 FR fish and 1255 NRL-NR (referred to as NR) fish.

## S4 Appendix Correlation between variables

Multicollinearity among continuous predictors was assessed using pairwise Pearson correlation coefficients (see Table S8). All pairwise correlations were low, with none exceeding 0.35, indicating the absence of strong linear relationships between variables. Based on this, all continuous predictors were retained for further analysis.

## S5 Appendix Justification of random effect

To verify if the random effect significantly improved model fit, we compared a mixed-effects model with a corresponding fixed-effects model. Including the random intercept for trial session ID substantially improved model performance (ΔBIC = 92.16). The likelihood ratio test confirmed that this effect was highly significant (*χ*^2^ = 99.37, df = 1, *p <* 0.001).

## S6 Appendix Effect of experimental variables

Spatial positioning relative to the stimulus (DSt and VA) played a significant role in determining whether a fish initiated an escape response. In the univariable analysis, distance to the stimulus (DSt) had a strong negative effect on response probability (*β* = *−*1.06 *±* 0.15, *p <* 0.001), explaining 15% of the variance (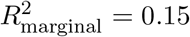; Table S2). This effect remained significant in the multivariable model (*β* = *−*0.98 *±* 0.19, *p <* 0.001; Table S3), and a log-transformation of DSt substantially improved model fit (ΔBIC = *−*7.14; Table S4). The binned data show a decline in the fraction of responses with increasing distance to the stimulus (Fig S3A). The angle at which the stimulus was viewed also influenced response probability. Viewing angle (VA) had a negative effect in both the univariable (*β* = *−*0.66 *±* 0.11, *p <* 0.001; Table S2) and multivariable models (*β* = *−*0.64 *±* 0.14, *p <* 0.001; Table S3). This trend is also present in the binned visualization, where the fraction of responses declines with increasing viewing angle (Fig S3B).

## S7 Appendix Alternatives for neighbor proximity

As an alternative to neighbor proximity (NPro), two other variables that represent social context were tested. The total angular area of the neighboring fish in the eye of the focal fish (NAS) and the total angular area in the side of the visual field that contained the stimulus (N). Both were calculated based on the size of the neighboring fish and the distance between the focal fish and the neighbors. As the sizes of the fish were prone to annotation errors due to their small size, the neighbor proximity (which did not depend on fish size) was the most reliable variable, and therefore used in the primary analysis.

When the two variables were tested in univariable analysis, N had a significant influence on the response probability (*β* = *−*0.47 *±* 0.15, *p <* 0.05). In the multivariable analysis, this effect became non-significant. This loss of significance likely reflects the confounding effect of a strong predictor in the model, specifically the distance to the stimulus, with which N has a weak correlation (*r* = 0.22). If fish were further from the stimulus, they had more neighbors in the visual hemisphere containing the stimulus. In both analyses, NAS was non-significant.

---

**Table S1.** Description of predictor variables used in the models. List of all experimental and ecological predictors, their abbreviations, and units.

| Variable | Symbol | Description | Units |
| --- | --- | --- | --- |
| Species | Sp | Species (*Bicolor Damselfish* vs. *Brown Chromis*) | Categorical |
| Orientation angle | OA | Orientation of fish relative to the stimulus | ° (0–180) |
| Viewing angle | VA | Angle at which the fish views the stimulus | ° (0–180) |
| Distance to coral | DCo | Distance to the closest point on the coral | *m* |
| Distance to stimulus | DSt | Distance to the center of the stimulus | *m* |
| Stimulus speed | LSp | Looming speed of the stimulus | $\text{m s}^{-1}$ |
| Trial event | TEv | Repetition of stimulus event in trial | Ordinal |
| Neighbor proximity | NPro | Summed squared reciprocal distance to all neighbors | $\text{m}^{-2}$ |

**Table S2.** Results of univariable GLMMs predicting escape probability. Each model included a single predictor and a random intercept for trial session. Estimates are shown with standard errors, p-values, and marginal R^2^ values.

| Variable | Estimate | Std. Error | p-value | $R^2_{\text{marginal}}$ |
| --- | --- | --- | --- | --- |
| Species (Sp) | −1.6570 | 0.3674 | $6.49 \times 10^{-6}$ | 0.1108 |
| Orientation angle (OA) | −0.0596 | 0.1207 | 0.622 | 0.0006 |
| Viewing angle (VA) | −0.6639 | 0.1053 | $2.83 \times 10^{-10}$ | 0.0663 |
| Distance to coral (DCo) | 0.6386 | 0.1178 | $5.85 \times 10^{-8}$ | 0.0634 |
| Distance to stimulus (DSt) | −1.0630 | 0.1449 | $2.22 \times 10^{-13}$ | 0.1523 |
| Stimulus speed (LSp) | 1.2550 | 0.1457 | $7.03 \times 10^{-18}$ | 0.1861 |
| Trial event (TEv) | −0.1810 | 0.1195 | 0.130 | 0.0053 |
| Neighbor proximity (NPro) | −0.6510 | 0.6515 | 0.318 | 0.0640 |

**Table S3.** Results of multivariable GLMM predicting escape probability. The model includes all predictors simultaneously, with random intercepts for trial session. Estimates, standard errors, z-values, and p-values are reported.

| Variable | Estimate | Std. Error | z-value | p-value |
| --- | --- | --- | --- | --- |
| Intercept | −3.6891 | 0.6443 | −5.726 | $1.03 \times 10^{-8}$ |
| Species (Sp) | −1.0994 | 0.4812 | −2.285 | 0.022 |
| Orientation angle (OA) | −0.0455 | 0.1446 | −0.315 | 0.753 |
| Viewing angle (VA) | −0.6385 | 0.1379 | −4.629 | $3.68 \times 10^{-6}$ |
| Distance to coral (DCo) | 0.1059 | 0.1723 | 0.615 | 0.539 |
| Distance to stimulus (DSt) | −0.9836 | 0.1904 | −5.165 | $2.40 \times 10^{-7}$ |
| Stimulus speed (LSp) | 1.3717 | 0.1680 | 8.164 | $3.24 \times 10^{-16}$ |
| Trial event (TEv) | −0.2589 | 0.1576 | −1.643 | 0.100 |
| Neighbor proximity (NPro) | −0.6228 | 0.7369 | −0.845 | 0.398 |

**Table S4.** Comparison of non-linear transformations for predictors in the context of the multivariable model. Comparison of BIC values for linear, log-transformed, and natural spline forms (2 or 3 *df*) of each predictor in the multivariable model. Best-fitting transformation per variable is indicated.

| Variable | BIC (linear) | BIC (log) | BIC (spline 2 df) | BIC (spline 3 df) | Best form |
| --- | --- | --- | --- | --- | --- |
| Orientation angle (OA) | 402.665 | 402.685 | 407.798 | 414.945 | Linear |
| Viewing angle (VA) | 402.665 | 406.729 | 406.898 | 413.451 | Linear |
| Distance to coral (DCo) | 407.587 | 402.665 | 408.084 | 411.766 | Log |
| Distance to stimulus (DSt) | 409.803 | 402.665 | 411.602 | 412.182 | Log |
| Stimulus speed (LSp) | 428.864 | 407.657 | 407.042 | 402.665 | Spline (3 df) |
| Trial event (TEv) | 402.665 | 405.192 | 404.945 | 401.328 | Spline (3 df) |
| Neighbor proximity (NPro) | 402.665 | 404.191 | 409.695 | 416.149 | Linear |

**Table S5.**
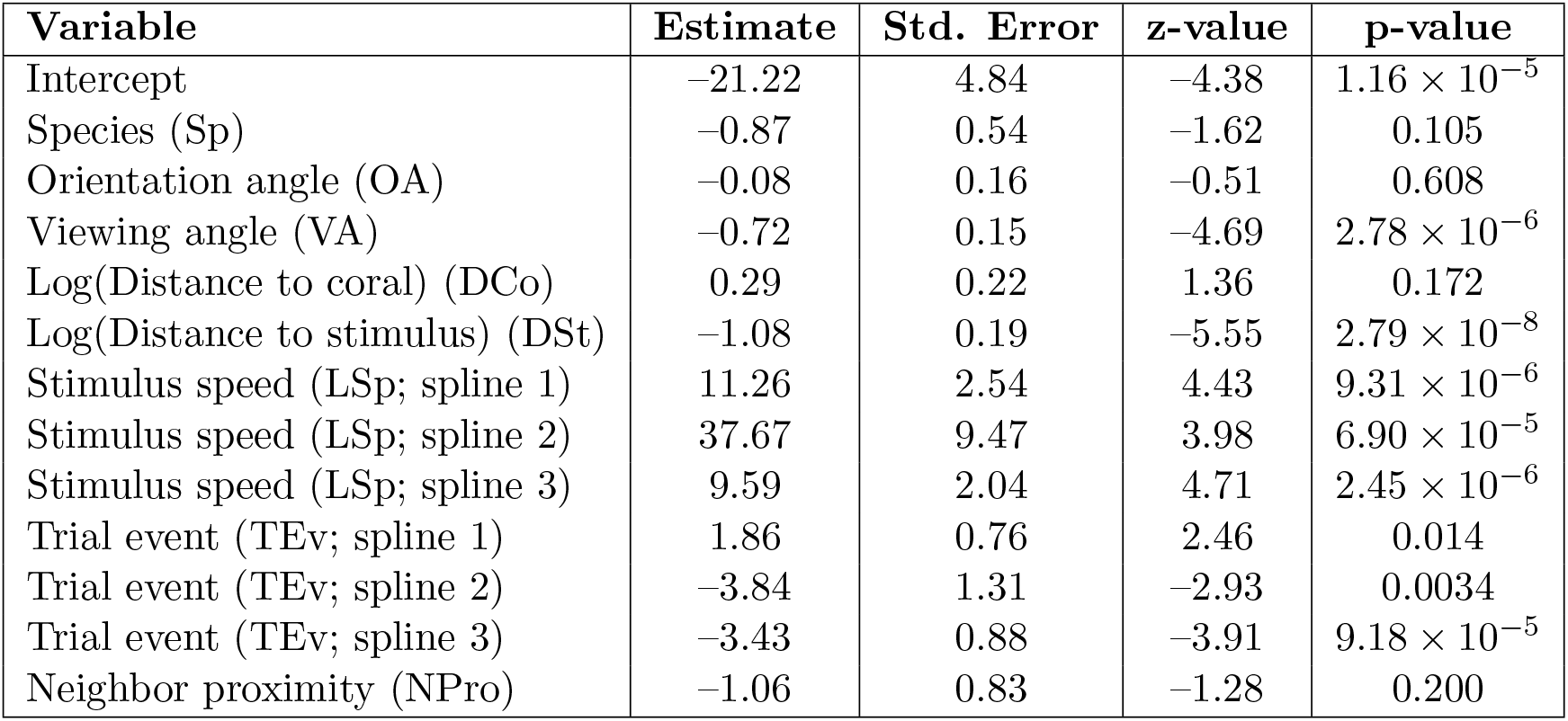
Final model results using natural spline terms for stimulus speed and trial order. Results from the best-fitting model, including spline terms for stimulus speed and trial event, and log-transformations of distance to coral and distance to stimulus. Estimates, standard errors, z-values, and p-values are provided.

**Table S6.** Effect of adding individual predictors to the species-only model. Comparison of AIC values and species coefficients when adding each predictor separately to a baseline species-only model. Reduction in the species coefficient indicates predictors that account for species differences.

| Predictor added | AIC | Species effect | SE | p-value |
| --- | --- | --- | --- | --- |
| None (baseline) | 546.8 | -1.66 | 0.367 | $6.5 \times 10^{-6}$ |
| Orientation angle (OA) | 548.6 | -1.66 | 0.368 | $6.5 \times 10^{-6}$ |
| Viewing angle (VA) | 513.3 | -1.59 | 0.386 | $3.7 \times 10^{-5}$ |
| Distance to coral (DCo; log) | 528.4 | -1.03 | 0.409 | 0.01 |
| Distance to stimulus (DSt; log) | 489.5 | -1.41 | 0.391 | $3.3 \times 10^{-4}$ |
| Stimulus speed (LSp; splines) | 429.7 | -1.71 | 0.410 | $3.0 \times 10^{-5}$ |
| Trial event (TEv; splines) | 544.0 | -1.66 | 0.370 | $7.7 \times 10^{-6}$ |
| Neighbor proximity (NPro) | 547.2 | -1.65 | 0.367 | $6.6 \times 10^{-6}$ |

**Table S7.** Ecologically relevant interaction effects, added to the multivariable model with LSp as natural splines. Ecologically relevant two-way interactions were tested with likelihood ratio tests.

| Variable 1 | Variable 2 | Effect Var 1 | Effect Var 2 | $\chi^2$ | Raw p-value | Adjusted p-value |
| --- | --- | --- | --- | --- | --- | --- |
| LSp | DCo | Splines (3 df) | Log | 9.13 | 0.0276 | 0.165 |
| LSp | Sp | Splines (3 df) | Categorical | 5.02 | 0.170 | 0.324 |
| LSp | NPro | Splines (3 df) | Linear | 4.78 | 0.189 | 0.324 |
| NPro | Sp | Linear | Categorical | 1.53 | 0.216 | 0.324 |
| DCo | Sp | Log | Categorical | 1.00 | 0.318 | 0.382 |
| NPro | DCo | Linear | Log | 0.56 | 0.454 | 0.454 |

**Table S8.** Correlation matrix of continuous predictors using Pearson correlation coefficients. Pearson correlation coefficients among continuous predictors. All correlations are *<* 0.35, indicating no strong collinearity.

| Variable | OA | VA | DCo | DSt | LSp | TEv | NPro |
| --- | --- | --- | --- | --- | --- | --- | --- |
| OA |  | 0.043 | −0.020 | 0.014 | 0.065 | 0.018 | 0.036 |
| VA |  |  | −0.293 | 0.305 | 0.036 | 0.023 | 0.021 |
| DCo |  |  |  | −0.277 | 0.071 | −0.004 | −0.073 |
| DSt |  |  |  |  | −0.015 | 0.030 | 0.006 |
| LSp |  |  |  |  |  | −0.026 | 0.023 |
| TEv |  |  |  |  |  |  | 0.028 |
| NPro |  |  |  |  |  |  |  |

## Notes

### Competing Interest Statement

The authors have declared no competing interest.

### Summary of Updates

Layout-issues, and addition of supplementary figures

